# Transmembrane Kinases are essential for plant development

**DOI:** 10.1101/2021.03.11.434919

**Authors:** Qiang Li, Jie Yang, Yi Zhang, Fen Wang, Mingzeng Chang, Tongda Xu, Shui Wang, Jun He

**Author notes:** Corresponding author: Jun He and S. W. These authors contributed equally to this work (Qiang Li, Jie Yang).

## Abstract

Transmembrane kinase family proteins (TMKs) have been implicated in regulating both auxin signaling and plant development. To obtain insights of the potential TMKs’ function in plant development, we regenerated new full sets of *tmk* mutants, and discovered many new phenotypes, such as defective organogenesis, smaller rosette leaves, fertility defects and selfing population defects in different combination of *tmk* mutants. Taken together, our results demonstrated that TMKs participated in multiple aspects of plant development, which provided a great reference for any future research.

**One-sentence summary:** Multiple *tmk* mutants analysis illustrated the key roles of TMKs in plant development.

## INTRODUCTION

Plant growth and development are regulated by different triggers like phytohormones or environments (Santner and Estelle, 2009, Csaba, 2015). Auxin, as the first identified phytohormone, is involved in nearly all aspects of plant lifecycle processes (Mockaitis and Estelle, 2008). The previous study suggested that auxin is required in the apical-basal axis formation in egg cells before fertilization (Mansfield et al., 1991, Laux and Jurgens, 1997), in the root apical meristem and shoot apical meristem maintenance (Sassi and Traas, 2015, Azizi et al., 2015, Della Rovere et al., 2013), in cotyledon initiation during the embryogenesis transition stage (Moller and Weijers, 2009) and in the formation of the lateral roots (Benkova et al., 2003, Dubrovsky et al., 2008).

Auxin signal was reported to be recognized and transduced by its binding protein the cell surface based (Auxin binding protein 1)ABP1 (Hertel et al., 1972, Dharmasiri et al., 2005, del Pozo et al., 2006, Jurado et al., 2008). ABP1 specifically binds with auxin with high affinity, but its developmental role is still unclear (Gao et al., 2015). TMK1 is identified as the docking protein of ABP1 at the cell surface, which mediates auxin activation of Rho GTPase signaling pathway (Klambt, 1990, Robert et al., 2010, Xu et al., 2010). TMK proteins are members of a subfamily of Leucine-rich repeat RLK family, which contains four family members. All four TMKs have the conserved LRR motif, the transmembrane region and the cytoplasmic kinase domain (Dai et al., 2013). The first identified member of them is TMK1 and it was characterized in the course of a chromosome walk to the ethylene response locus, and further proved as a kinase for serine and threonine by in vitro kinase assay (Chang et al., 1992). The kinase exhibited greater autophosphorylating activity with Mn^2+^ than with Mg^2+^(Schaller and Bleecker, 1993). Recent work demonstrates that auxin accumulation at the concave side of the apical hook stimulates TMK1 cleavage followed by its nuclear translocation to regulate gene transcription via stabilizing IAA32 and IAA34, two non-canonical Aux/IAA proteins lacking domain* which is required for interaction with the TIR receptors (Cao et al., 2019). TMK4 was first cloned as BAK1-Associating Receptor-Like Kinase 1 (BARK1) and reported to regulate BR-mediated plant development including lateral roots via auxin regulation (Kim et al., 2013). Moreover, TMK4 was reported to interact with and phosphorylates TAA1 at Thr101 site, which negatively regulates auxin biosynthesis that is essential for the regulation of the local auxin concentration in plants (Wang et al., 2020). And also TMK1/TMK4-mediated phosphorylation and activation of MKK4/MKK5, followed by the activation of MPK3/MPK6, are involved in cell division during lateral root (LR) development (Huang et al., 2019). All these indicate the multiple functions of TMKs at least in auxin signaling.

The T-DNA insertion mutants of these four TMKs were isolated previously (Dai et al., 2013, Schwachtje et al., 2012). And no obvious developmental phenotype was observed in the *tmk* single mutants. But in the *tmk1/4, tmk1/3/4* and *tmk1/2/3/4* mutants, the development of the root, hypocotyl and stamen filament was strongly reduced due to both cell division and elongation defects (Dai et al., 2013). The *tmk1/2/3/4* quadruple mutant is sterile (Schwachtje et al., 2012, Dai et al., 2013). Moreover, the *tmk* mutants showed less sensitivity to exogenous auxin treatment in primary root growth, lateral root initiation and pavement cell interdigitation (Dai et al., 2013, Xu et al., 2014). The comprehensive phenotypes in multiple *tmk* mutants suggest the essential roles of TMKs in plant development, which needs to be further carefully investigated. Here, by regenerating different combinations of multiple *tmk* mutants in pure Col-0 background, we further extended our understanding of the TMKs function in regulating plant development, including vegetative development and fertilization processes. Taken together, our results demonstrated that TMKs modulate organ developmental programming and adaptive growth to the environmental stimulus, which is essential for *Arabidopsis* development.

## RESULTS

### Novel *tmk* T-DNA insertion mutants with pure Col-0 background

The phenotypes of *Arabidopsis* varied excessively among different ecotypes (Schwachtje et al., 2012). Although the important roles of TMKs in auxin signaling, previous *tmks* with mixed ecotypes might be debated in the whole TMK and auxin research society. There are total 4 members of TMKs subfamily which locate at different chromosomes in *Arabidopsis* genome (Figure 1A). To study the functions of these 4 members of TMKs subfamily in *Arabidopsis*, we collected and identified each *TMKs* single knock-out T-DNA mutant in Col-0 background from publicly available resources. The *tmk1-/-*(*tmk1*) single mutant was the same as before because it is Col-0 background already (Dai et al., 2013). Novel *tmk2, tmk3* and *tmk4* knockout mutants were identified and the T-DNA insertion is located at positions +1550 bp, +2106 bp, +137 bp, +2559 bp relative to the start codons of their corresponding genes, respectively (Figure 1B). The expression of each *TMK* in the corresponding individual *tmk* single mutant was confirmed by genotyping and quantitative real-time RT-PCR (qRT-PCR) (Figure1D and 1E). To generate novel *tmk* mutant combinations, the *tmk1* single mutant was crossed with the novel *tmk2-/- (tmk2)* single mutant and, meanwhile, the novel *tmk3-/- (tmk3)* single mutant was crossed with the novel *tmk4-/-(tmk4)* single mutant, respectively. The F1 generation lines *tmk1+/-tmk2+/-* and *tmk3+/-tmk4+/-* were then crossed with each other. In the next generation, the heterozygous line *tmk1+/-tmk2+/-tmk3+/-tmk4+/-* was obtained by genotyping. The seeds of this line were collected to further screen candidates for double, triple, and quadruple *tmk* mutants. Finally, different combinations of novel *tmk* mutants, including 4 *tmk* single mutants, 6 *tmk* double mutants, 4 *tmk* triple mutants and one *tmk* quadruple mutant were obtained.

**Figure 1.**
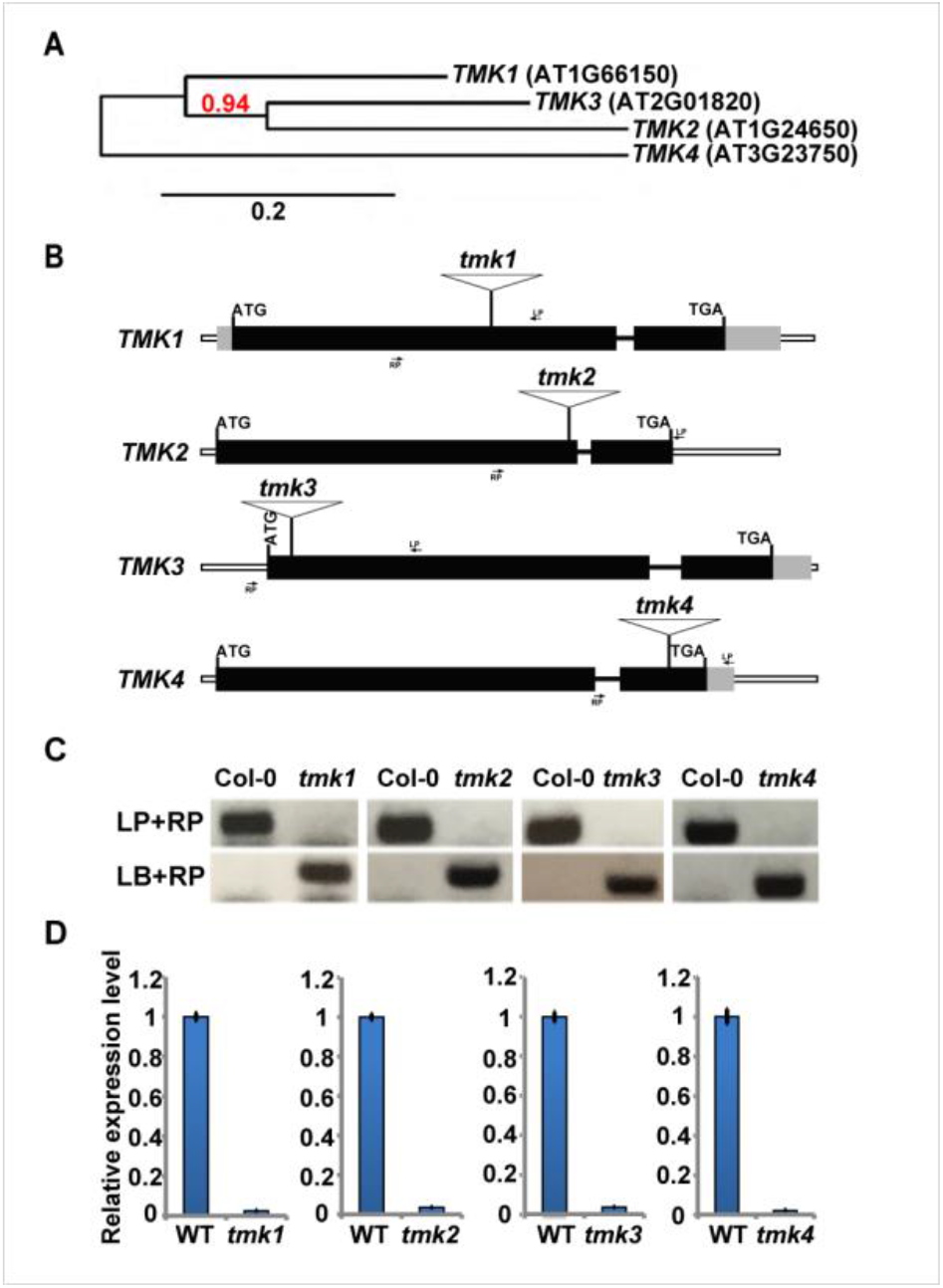
Generation of novel mutant combinations among *TMKs*. **(A)** The phylogenetic tree of *TMKs* in *Arabidopsis* genome predicted by MEGA 5. **(B)** Identification of the T-DNA insertion sites of the *tmk* mutants. The black boxes, gray boxes, lines, and long boxes indicate the exons, untranslated regions, introns, and interval regions, respectively The locations of LB and RP primer pair are indicated by arrows. **(C)** Genotyping of 4 *tmk* single mutants. No bands show up in *tmk1*, novel *tmk2, tmk3*, and *tmk4* single mutants as compared to Col-0 in the gene specific PCR groups (the upper panel). No bands present in Col-0, but a band is shown in *tmk1* and novel *tmk2, tmk3*, and *tmk4* single mutant in the T-DNA specific PCR group (the lower panel). **(D)** Gene expression of *TMKs* in their corresponding mutants by using qRT-PCR. The transcripts of *TMK1, TMK2, TMK3*, and *TMK4* are almost undetectable in the *tmk1* and novel *tmk2, tmk3, tmk4* single mutants, respectively. *UBQ1* was applied as an internal control. Error bars are SD.

### TMKs are required for vegetative development in *Arabidopsis*

We further examined carefully whether there are any abnormal vegetative developmental phenotypes in these new *tmk* mutants. We focused on 4 developmental characters, the biggest rosette leaf (BRL) at the juvenile stage and the primary inflorescence height (PIH) at the adult stage, the primary root length (PRL) and etiolated hypocotyl length. All *tmk* single mutants showed no obvious defects (Figure 2), except *tmk4* single mutant which exhibited obvious growth defects in BRL (around 22.3% reduction) (Figure 2A and 2B) and PRL (around 26% reduction) in PRL comparing to Col-0 wild type (Figure 2E and 2F). In contrast, severe developmental defects were observed in *tmk1-/-tmk4-/-* (*tmk1/4*) double, *tmk1-/-tmk2-/-tmk4-/-* (*tmk1/2/4*) and *tmk1-/-tmk3-/-tmk4-/-* (*tmk1/3/4*) triple and *tmk1-/-tmk2-/-tmk3-/-tmk4-/- (tmk1/2/3/4)* quadruple mutants. The BRL in *tmk1/4* double mutant and *tmk1/2/3/4* quadruple mutant was around 38% and 21% of the Col-0, respectively (Figure 2A and 2B). The PIH in *tmk1/4* double mutants was 59% of the Col-0, and only 12% that of the Col-0 in the *tmk1/2/3/4* quadruple mutants (Figure 2C and 2D). The *tmk1/2/4* and *tmk1/3/4* triple mutants displayed slightly severe phenotype at BRL and PIH compared with *tmk1/4* double mutants (Figure 2A, 2B, 2C, and 2D). The PRL in *tmk1/4, tmk1/2/4*, and *tmk1/3/4* mutants was around 1/3 that of the Col-0, whereas about 1/6 that of the Col-0 in the *tmk1/2/3/4* quadruple mutants was observed at the early stage of seedling development (Figure 2E and 2F). For the hypocotyl length, *tmk1/2/3/4* was shortest (how much reduction) followed by *tmk1/4* (70% reduction) compared to Col-0. Furthermore, we confirmed that the *tmk1/4* complementation line (complemented by native promoter driven *gTMK1*,see experimental procedures) rescued the hypocotyl phenotype (data not shown). Taken together, *tmk1//2/3/4* shows the most severe growth defect phenotypes followed by *tmk1/2/4 or tmk1/3/4* and *tmk1/4*. Therefore, we concentrated on these mutants, especially *tmk1/4* and *tmk1/2/3/4*, for further study.

**Figure 2.**
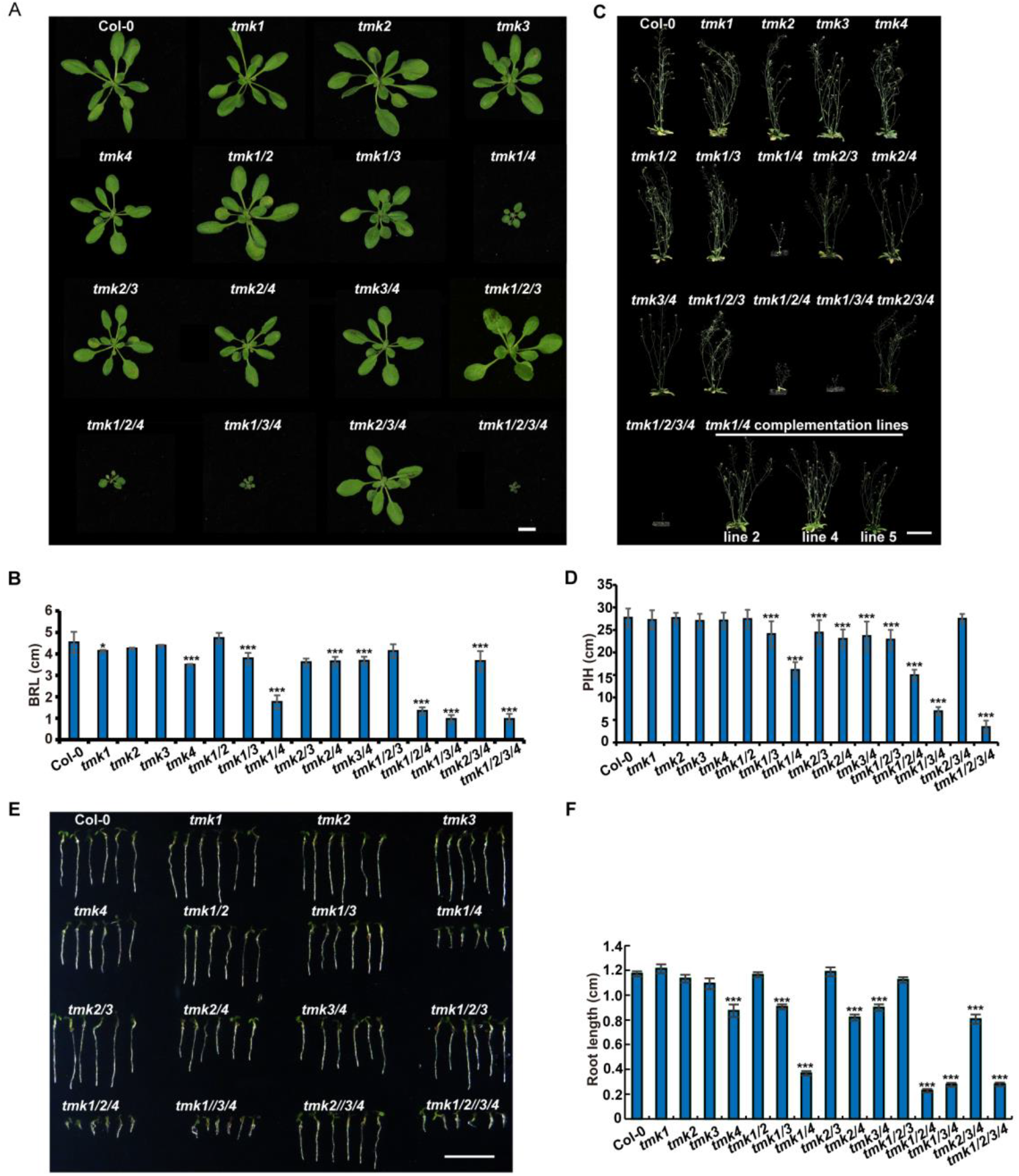
The phenotype characterization of the novel *tmk* mutants and the complementation lines. **(A)**The 24-dag-old (grown in a growth chamber for 10 days and transferred to soil and grown in the greenhouse for another 14 days) plants of novel *tmk* mutants and Col-0. Bar, 10 cm. **(B)** Quantitative analysis of rosette leaf size of novel *tmk* mutants. **(C)** The 45-dag-old (10-dag-old seedlings transferred to soil and grown for another 35 days) plants of novel *tmk* mutants, Col-0, and the *tmk1/4* complementation lines (*tmk1/4:gTMK1*, see experimental procedures). Bar, 5 cm. **(D)**Quantitative analysis of primary inflorescence height (PIH). Shown are average ± SD (n=30). **(E)** The 4-dag-old seedlings of novel *tmk* mutants and the Col-0. **(F)** Quantification of root length. Shown are average ± SD (n=30). Note: t-test for comparison between *tmk* mutants and Col-0. Error bars denote ±SD. “*”, “**”, and “***” indicate p<0.05, p<0.01, and p<0.001, respectively.

In previous *tmk1+/-tmk2-/-tmk3-/-tmk4-/-* (*tmk1*+/-*tmk2/3/4)* heterozygous lines, the single cotyledon phenotype was observed (Xu et al., 2014). Interestingly, besides single cotyledon, we also observed triple and swollen cotyledons in our novel *tmk1-/-tmk2-/-tmk3-/-tmk4+/-* (*tmk1/2/3;tmk4+/-*) mutants (Figure 4A and 1B). *The tmk1/2/3* triple mutants displayed cotyledon-shaping defects as well, although with an even lower frequency (about 0.26%) (Figure 4C). Furthermore, the rootless phenotype coupled with triple cotyledons in novel *tmk1/2/3/4* quadruple mutants were also identified (Figure 4B).

Taken together, our data suggested that functions of TMKs in *Arabidopsis* vegetative growth from seedling stage to adult stage are essential and largely overlapped. In addition, roles of TMK4 were irreplaceable by the other 3 TMKs during *Arabidopsis* vegetative growth. Since *tmk1//2/3/4* showed the most severe growth defect phenotypes followed by *tmk1/2/4 or tmk1/3/4* and *tmk1/4*. Therefore, we focused on these mutants, especially *tmk1/4* and *tmk1/2/3/4*, for further study.

### TMKs are required for *Arabidopsis* reproductive growth

Next, we examined whether there are any abnormal phenotypes at reproductive growth stages in these new *tmk* mutants. We found that the *tmk1/2/3/4* mutant showed the shortest inflorescence shoot and silique phenotype followed by *tmk1/4* mutant but no obvious phenotypes were observed in *tmk* single mutants (Figure 3A and 3B). In agreement with the reduced silique length, the mature seed number was dramatically decreased, in *tmk1/4, tmk1/2/4*, and *tmk2/3/4* mutants, to 18.8%, 14.5%, and 41.6% that of wild type, respectively (Figure 3C). Strikingly, there are no matured seeds produced in siliques of *tmk1/3/4* and *tmk1/2/3/4* mutants (Figure 3C), which implied an essential role of TMKs in the fertilization process or embryo development. To further investigate this hypothesis, whole flower organ of *tmks* mutants were observed firstly, among which *tmk1/2/3/4* displays obvious size reduction followed by *tmk1/4* combinations (*tmk1/3/4, tmk1/2/4*, and *tmk1/4*) and *tmk2/3/4* (Figure 3D). We further split out the flowers of these mutants and found the petal, sepal, pistil, and stamen are all reduced in size/length, especially in the *tmk1/2/3/4* quadruple mutant, although no shapes and numbers seem to be defective (Figure 3 E-G). When peeling off the silique, we found there are totally aborted seeds that seem to arise from fertility in the aforementioned *tmk* mutants (Figure 3H). From this part of the view, *tmk1*/*2/3/4* quadruple mutants were completely infertile, followed by *tmk1/4* and *tmk2/3/4* mutants, whereas no fertility defects in single *tmk* mutants (Figure 3H). Occasionally, it was interesting to identify that all the stamens in novel *tmk1/2/3/4* quadruple mutants were shorter than the stigma (Figure 3F and 3G). Thus, to study the infertility phenotype of *tmk* mutants caused by the stamen or the pistil, the pollen of the *tmk1/2/3/4* quadruple mutant was smeared on the stigma of Col-0, and then (3 weeks later) the siliques were split out and only 3 to 5 mature seeds could be obtained in one silique on average (Figure 3I, the left panel, indicated by red arrows) suggesting the anther from *tmk1/2/3/4* quadruple mutant has extremely severe fertility defects. No mature seeds could be observed in the siliques no matter the smeared pollen was from Col-0 or the novel *tmk1/2/3/4* quadruple mutant (Figure 3I, the middle and right panels, indicated by red arrows) suggesting the pistil from *tmk1/2/3/4* loses the fertility thoroughly. Based on these results, the sterility of *tmk1/2/3/4* quadruple mutants might be caused by defects in both stamen elongation and pistil development. Thus, TMKs might play previously unknown roles in the regulation of stamen elongation, ovule and pollen development, and seed maturation. In addition, according to the very low ratio (7.69% + 0.38% = 8.07%, which is far less than the theoretical ratio 25%; Figure 4B) of homozygous *tmk1/2/3; TMK4-/-* from the *tmk1/2/3; TMK4+/-* parental generation, it is noteworthy that TMKs must also play a role in embryo development.

**Figure 3.**
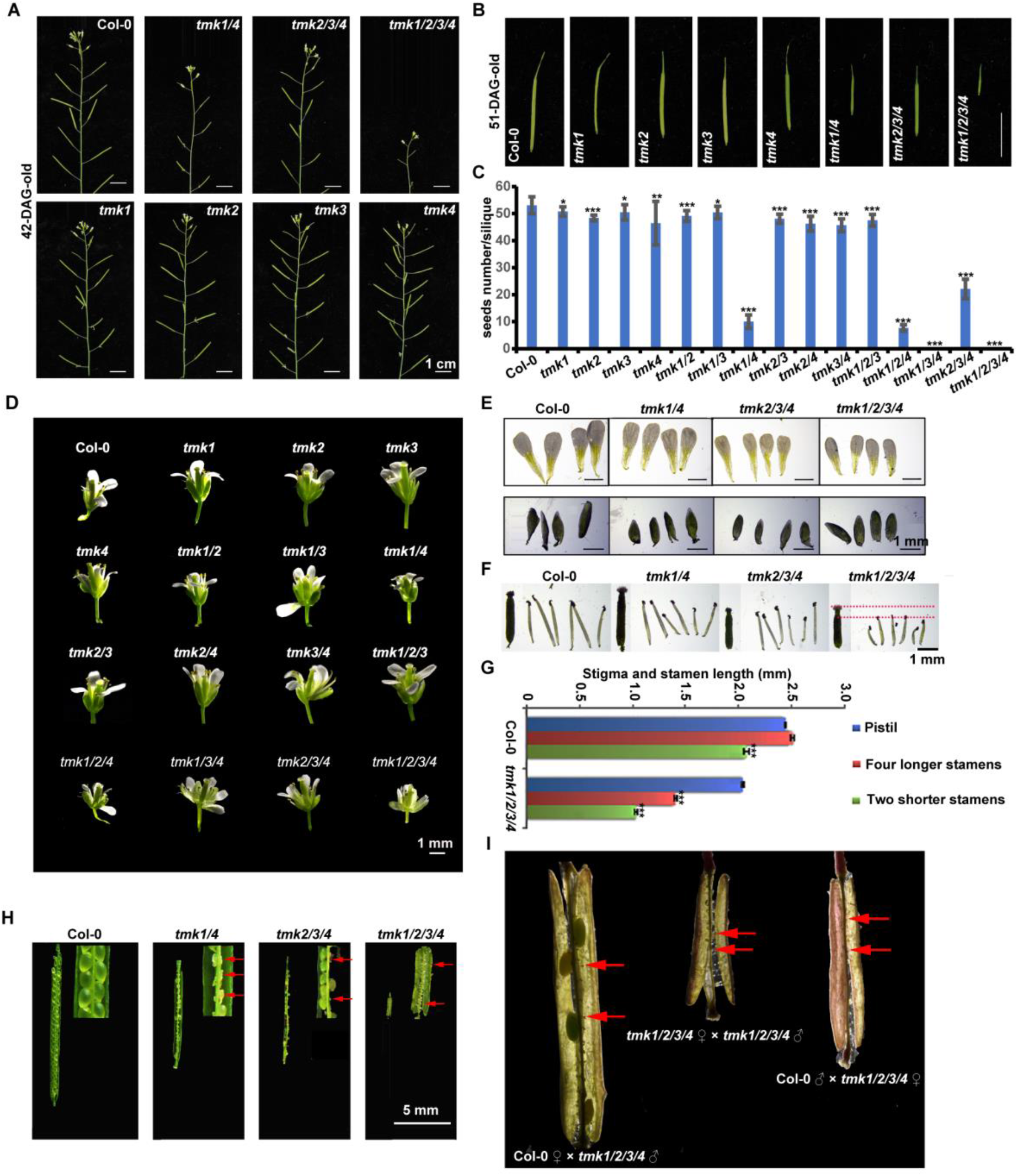
The novel *tmk* mutants displayed sexual growth defects. **(A)**The shoots of 42-dag-old Col-0, *tmk* single mutants, *tmk1/4, tmk2/3/4*, and *tmk1/2/3/4* mutants. Bars = 1 cm. **(B)** The siliques of 51-dag-old Col-0, *tmk* single mutants, *tmk1/4, tmk2/3/4* and *tmk1/2/3/4* mutants. Bar, 1 cm. **(C)** Quantitative analysis of the seeds number per silique in Col-0, *tmk* single mutants, *tmk1/4, tmk2/3/4* and *tmk1/2/3/4* mutants. **(D)** The intact flower of novel *tmk* mutants and Col-0 (45-d-old). Bar, 1 mm. **(E)** The sepals and petals comparison in 45-dag-old Col-0 and *tmk1/4, tmk2/3/4* and *tmk1/2/3/4* mutants. Bar, 1 mm. **(F)** The comparison of the stamens and stigma length of the novel *tmk1/4, tmk2/3/4*, and *tmk1/2/3/4* mutants and Col-0. The shortened length of stamen compared with stigma in *tmk1/2/3/4* mutants was noted by red lines. Bar, 1 mm. **(G)** Quantitative analysis of stigma and stamen length in Col-0 and *tmk1/2/3/4* mutant. **(H)** Aborted seeds maturation in the novel *tmk* mutants. Bar, 5 mm. **(I)** The emasculated Col-0 flower that was pollinated with novel *tmk1/2/3/4* mutant pollen developed only 3-5 mature seeds (the left panel); the emasculated novel *tmk1/2/3/4* mutant flower that was pollinated with its own pollen and Col-0 pollen developed no seeds (the middle and right panels). Note: t-test for comparison between *tmk* mutants and Col-0. Error bars denote ±SD. “*”, “**”, and “***” indicate p<0.05, p<0.01, and p<0.001, respectively.

**Figure 4.**
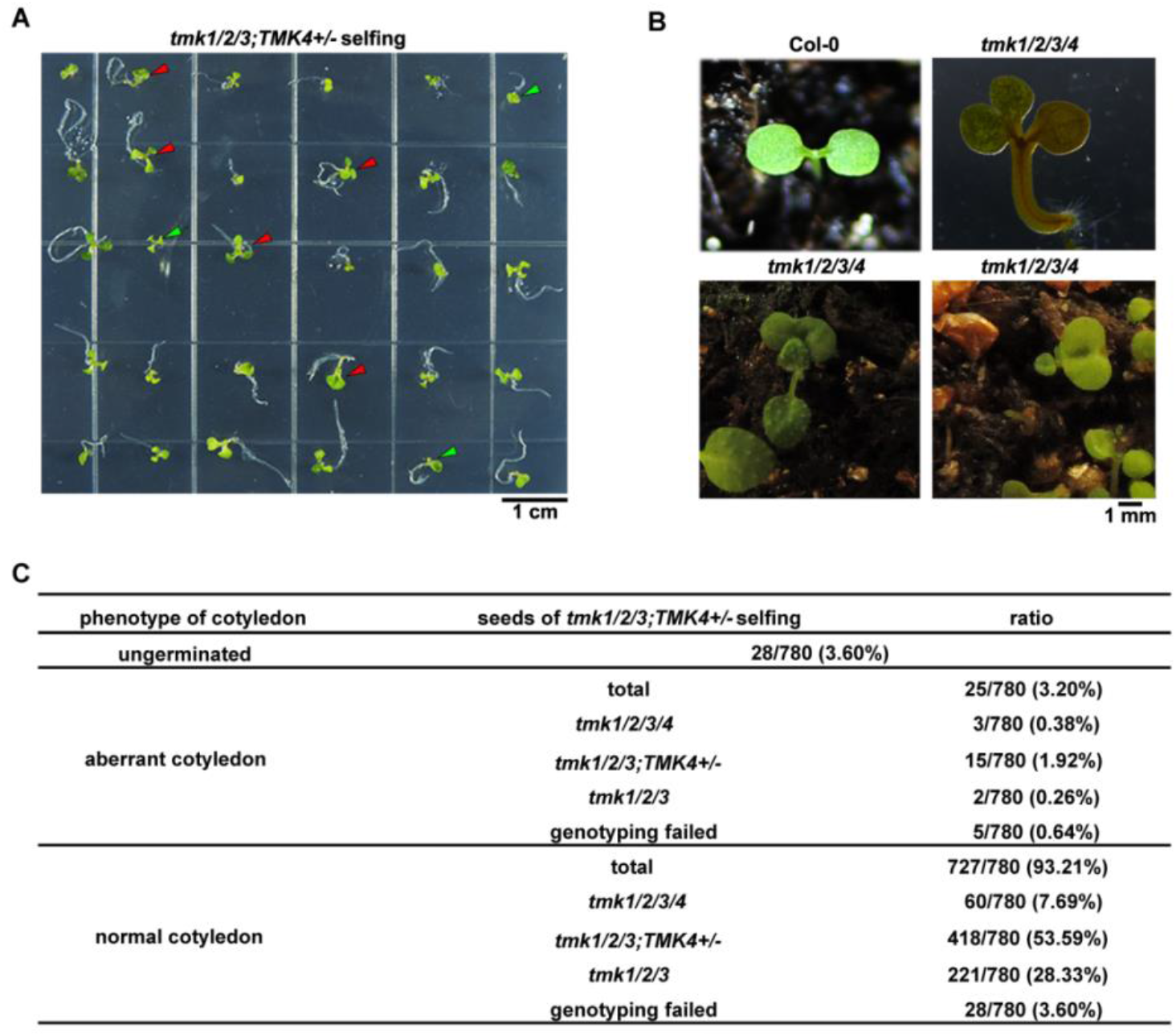
Phenotype analysis of *tmk123;TMK4+/-* selfing populations. **(A)**Single, triple and swollen cotyledons of novel *tmk* mutants from the progenies of *tmk1/2/3;tmk4+/-* selfing. Among the progenies of the *tmk1/2/3;tmk4+/-* selfing, the triple cotyledons with long root (shown by red arrows), single cotyledon with short root (shown by green arrows), and inflated cotyledons (shown by blue arrows) were observed. Bar, 1 cm. **(B)** Enlarged phenotype of the abnormal cotyledons of *tmk1/2/3/4* mutants grown in soil. Col-0 was used as a control. Bar, 1 mm. **(C)** Quantitative analysis of the abnormal cotyledon ratio of progenies of *tmk1/2/3;tmk4+/-* selfing. In 780 seedlings of *tmk123;tmk4+/-* selfing progenies, 3.60% seeds failed to germinate, while 3.20% showed abnormal cotyledon phenotype, which includes 0.38% *tmk123;tmk4-/-*, 1.92% *tmk123;tmk4+/-*, and 0.26% *tmk123;tmk4+/+* (Chi-square test, p<0.01).

## DISCUSSIONS

Here, our findings present several important implications. First, our results confirmed functionally overlapping roles of the TMKs and pointed out the unique and irreplaceable status of TMK4 among four TMKs. Second, phenotype characterization of novel *tmk* mutant combinations expanded the roles of TMKs from fundamental growth to adapted growth in *Arabidopsis*. Third, our data provided genetic evidence more accurately and comprehensively for future studies of TMKs family. All these findings here render roles of TMKs to more diverse and more general aspects of *Arabidopsis* developments. Here we applied multiple model systems at different tissue/organ levels to investigate roles of TMKs in *Arabidopsis* development. Future studies are required for deciphering the mechanisms of TMKs mediated multiple developmental processes at the molecular and biochemical cellular levels.

### Characterizations of novel *tmk* mutants reveal TMKs orchestrate many fundamental aspects of *Arabidopsis* growth and development

In this paper, we generated and comprehensively studied novel *tmk* combination mutants with pure ecotype background. We characterized the loss function of *TMKs* as dampens both vegetative growth and reproductive growth including primary roots, hypocotyls, rosette leaves, inflorescence, flower organs, siliques, and seeds maturation. These results are greatly consistent with Bleecker’s group, who showed severe growth and development defects of *tmk1;tmk2;tmk3;tmk4* (they generated) quadruple mutant in roots, hypocotyls, leaves, and stamen filaments caused by hampered cell growth and proliferation, although the T-DNA insertion mutants of the four TMKs contains ecotype mixture (Dai et al., 2013). Similarly, among all the novel *tmk* mutant combinations, the novel *tmk1/4* double, *tmk1/2/4* and *tmk1/3/4* triple and *tmk1/2/3/4* quadruple mutants displayed especially severe defects in almost all examined development processes of diverse *Arabidopsis* organs, suggesting the outstanding roles of TMK1 and TMK4 among the four functional redundant TMKs in *Arabidopsis*. Unlike the previous *tmk4* single mutant, here we discovered obvious defects of novel *tmk4* single mutant in primary rosette leaf expansion (Figure 2A and 2B) and root elongation (Figure 2E and 2F). Novel *tmk1/2/3/4* displayed significantly enhanced defects in both primary inflorescence and pavement cells (PCs) interdigitation (Figure 2C and 2D) as compared to the previous one (Xu et al., 2014, Dai et al., 2013). These results emphasized the unique role of TMK4 among four TMKs in the vegetative growth of *Arabidopsis* and indicated that the previous mutants with mixed ecotypes might cover the phenotype defects both at the organic level and cellular level during *Arabidopsis* vegetative growth. In the reproductive growth of *Arabidopsis*, the phenotype defects covered by the ecotype mixture seem to be much stronger, although fertility in most novel *tmk* mutants was similar to the Col-0 and the novel *tmk1*/*2/3/4* quadruple mutants were also completely infertile like previous ones (Dai et al., 2013) (Figure 3). More severe infertility showed up in novel *tmk1/4* double mutants as compared to previous *tmk1/4* mutants: only 18.8% seeds per silique as compared to wild type were matured successfully in the silique of novel *tmk1/4* double mutants, rather than 30.7% that in the previous *tmk1/4* double mutants (Figure 3C; Dai et al., 2013). What’s more, the novel *tmk2/3/4* triple mutant also displayed severe infertility while the previous one didn’t show visible fertility defects. Finally, both severe fertility defects of stamen and completely fertility of pistil result in the infertility phenotype of *tmk1/2/3/4* mutant (Figure 3I). More interestingly, both the low homozygous ratio in progenies of *tmk1/2/3;tmk4+/-* and aborted seeds growth observed in *tmk1/4* and *tmk2/3/4* hint the embryo development defects also exist as confirmed in a previous paper by using previous *tmk1/2/3/4* mutant (Xu et al., 2014). Moreover, the ratio of abnormal cotyledon morphology in progenies of *tmk1/2/3;tmk4+/-* (3.2%) was lower than that of *tmk1-/+tmk234* (17.1%) (Figure 4C) (Xu et al., 2014). It is likely that the embryo development and phenotype in the novel *tmk* quadruple mutants must be easily affected by growth conditions because the homozygous *tmk1/2/3/4* ratio is 8.08% (Figure 4C), much lower than the 25% separation ratio, and also we can’t exclude the reason from ecotype mixture. These findings expanded the roles of TMKs to more detailed and deeper aspects of reproductive growth in *Arabidopsis*. Apart from *tmk1/2/3/4* quadruple mutant, PCs of all the other *tmk* mutant combinations are firstly examined here. We found that novel *tmk1/2/3/4* quadruple mutant displayed similar PCs interdigitation defects as reported (Figure 5) (Xu et al., 2014). Taken together, although the *tmk* mutants possess similar growth and development defects which match that of the previous one to a large content, several phenotypes observed here differed from previous ones which still cannot exclude the ecotype mixture problems. Therefore, the new mutants could be better research materials for future study.

**Figure 5.**
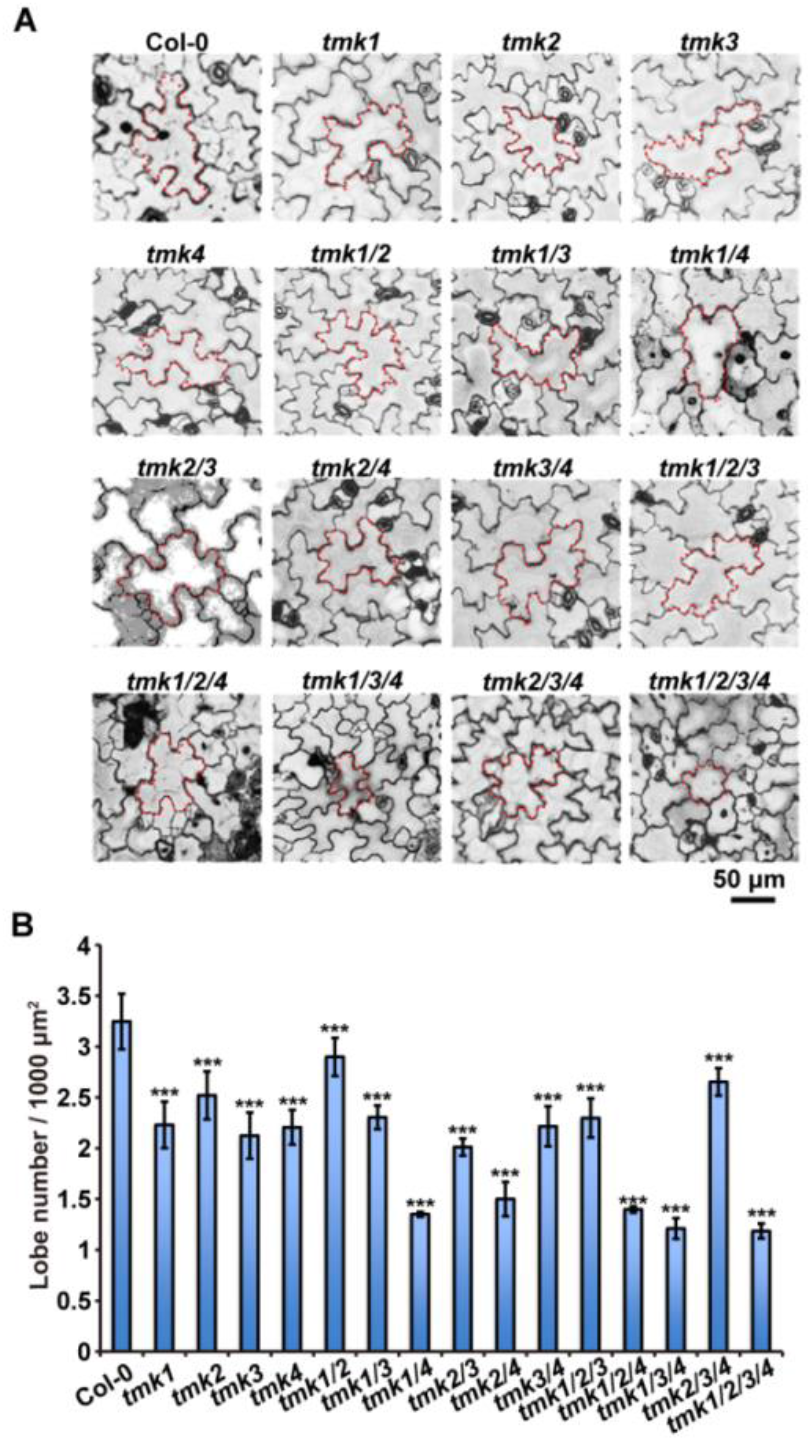
Pavement cell phenotype in Col-0 and the *tmk* combination mutants. **(A)**The pavement cell phenotype of Col-0 and all the generated *tmk* mutants. Bar, 50 μm. **(B)** The lobe number per μm^2^*1000 of Col-0 and all the generated *tmk* mutants. Note: t-test for comparison between *tmk* mutants and Col-0. Error bars denote ±SD. “*”, “**”, and “***” indicate p<0.05, p<0.01, and p<0.001, respectively.

### Function specificity of each TMK

By using promoter:GUS reporter system and qRT-PCR analyses, *TMK1, TMK3*, and *TMK4* were demonstrated to be widely expressed in all organs tested including roots, stems, leaves, flowers, and siliques, whereas *TMK2* expression was in trace amounts only detected in flowers and siliques of wild type *Arabidopsis* (Dai et al., 2013). We confirmed the knock-out *tmk1* and *tmk4* mutants through checking transcripts abundance by qRT-PCR (Figure 1). The relatively more important roles of TMK1 and TMK4 may contribute to the more prevalent expression patterns among organs in *Arabidopsis*. Substantial serious defects of novel *tmk1/4* and novel *tmk1/2/3/4* mutants are also consistent with the particular expression pattern of each TMK. TMK4 itself also plays roles in growth relate phenotypes such as rosette leaf size and primary root elongation (Figure 1A, 1B, 1E, and 1F) suggesting its dominant and specific function in these tissues. To our surprise, we also discovered the novel *tmk2/3/4* triple mutant showed reduced fertility (Figure 3G), which indicates that the roles of TMK2, TMK3, and TMK4 in controlling sexual growth. In the future, the expression pattern data with higher resolution should be obtained and examined carefully to separate the function of each TMK. What’s more, the expression level of each TMK in different *tmk* mutant combinations should be investigated simultaneously. And it’s worthy to note that apart from the T-DNA insertion lines, the other TMKs knock down or knock out lines are still necessarily going to be constructed by other technologies. As described above, TMK4 is also named as BRI1-ASSOCIATED RECEPTOR KINASE1 (BAK1)-associating receptor-like kinase 1 due to its binding to BAK1 (Kim et al., 2013). Further study in auxin-brassinosteroid crosstalk may consider TMK4 as the possible link component.

## EXPERIMENTAL PROCEDURES

### Plant Materials and Growth Condition

*Arabidopsis thaliana* seeds were surface-sterilized in 70% (v/v) ethanol for 10 minutes, and rinsed with sterile water 3 to 5 times. After stratified at 4°C for 2 to 5 days in the dark, seeds were placed on 1/2 Murashige and Skoog (MS) medium supplemented with 1% (w/v) sucrose and 0.8% (w/v) agar for 2 weeks in a growth chamber and then 2-week-old seedlings were transferred to soil for further growth in greenhouse. The growth condition of both the growth chamber and the greenhouse is at 22°C with a 16-h light/8-h dark cycle at 75 μmol m^-2^ s^-1^ and 65% humidity.

The T-DNA insertion lines of both *tmk1-/-* (SALK_016360) and *tmk2-/-* (SAIL_1242_H07) used here are previously reported single mutants with pure Columbia-0 (Col-0) ecotype. The Col-0 background *tmk3-/-* (SALK_129759) and *tmk4-/-* (GABI_191D02) single mutants were obtained from the Nottingham *Arabidopsis* Stock Centre (NASC; http://*Arabidopsis*.info). The transgenic lines *DR5p::GUS* and *DR5p::GFP* with Col-0 ecotype were kindly supported by Shingo Nagawa. *DR5p::GFP/GUS* with *tmk1/4* background (*DR5p::GFP /tmk1/4*) was generated from *DR5p::GFP/GUS* crossing with *tmk1/4* mutant. For three *tmk1/4* complementation lines driven by TMK1 promoter, *tmk1/4:gTMK1-2, tmk1/4:gTMK1-4*, and *tmk1/4:gTMK1-5*, please see the complementation assay.

### Identification of T-DNA Insertion Mutants

According to previous reports, genotype specific PCR was used for identifying T-DNA insertion sites of these Col-0 background *tmk* mutants. Generally, LB and RP primer set was used to identify T-DNA insertion while LP and RP primer set was used to confirm homozygous insertion. For the primer used, For LB primers, LBb1.3, LB1, and GABI-o8409-LB primer were used for “SALK”, “SAIL” and “GABI” mutants, respectively. The corresponding LP and RP primer sets were designed at the website (http://signal.salk.edu/tdnaprimers.2.html). For primer sequence, please see Supplemental Table 1. Procedure for PCR amplification by using ExTaq polymerase (Takara): 94°C 2 min; 35 cycles of 94°C 20 s, 62°C 30s, and 72°C 45s; 72°C 5 min.

### Complementation Assay

To generate the *tmk1/4:gTMK1* complementation line, 3.5 kb *TMK1* promoter driven genomic sequence plus 1 kb 3’-UTR of *TMK1* were PCR amplified and inserted into pDONR-Zeo vector to construct pDONR-gTMK1, FLAG tag was inserted into 3’-end of gTMK1 (pDONR-gTMK1-FLAG) and subsequently, gTMK1-FLAG was transferred into pGWB501 destination vector by LR reaction. The recombinant vector gTMK1-FLAG-pGWB501 was introduced into the *tmk1/4* mutant plants via *Agrobacterium tumefaciens*-mediated transformation (Clough and Bent, 1998). Transgenic plants were screened by 1/2 MS plates (1% sucrose) with 25 µg/ml of Hygromycin B. Finally, three T3 homozygous *tmk1/4* complementation lines driven by TMK1 promoter, *tmk1/4:gTMK1-2, tmk1/4:gTMK1-4*, and *tmk1/4:gTMK1-5*, were obtained (Figure 2C: line 2, line 4, and line 5).

### RNA Isolation, cDNA Synthesis and Quantitative Real-time RT-PCR (qRT-PCR)

5-dag (day after germination)-old seedlings of *tmk* mutants grown in the growth chamber were frozen and ground by liquid nitrogen. RNA was isolated by Trizol RNA isolation kit (Invitrogen). After RNA isolation, reverse transcription was performed using cDNA synthesis superMix (TransGen Biotech). Actually, total 500 ng to 1 μg RNA was added in a total reaction volume of 20 μL. Reverse transcribed cDNA was diluted to 10 times with ddH_2_O before quantitative real-time RT-PCR. For qRT-PCR, 2.5 μM forward and reverse primer set, 3 μL cDNA template and 4 μL iQ SYBR green supermix (Bio-Rad) in a total reaction volume of 12 μL. The reaction was detected by using CFX96™ Real-Time System (Bio-Rad). *TUB4 (AT5G44340)* was used as a control gene. For primer used to detect the *TMK* gene expression level, please see Supplemental Table 1.

### Phenotypic Analysis

Actually, “10+14” plants (10-dag-old seedlings grown in a growth chamber, transferred in soil and grown in the greenhouse for another 14 days) were used for the analysis of the diameter of the biggest rosette leaf (BRL). For measurement of the primary inflorescence height (PIH), “10+35” plants were used. For detection of the primary root length, 4-dag-old seedlings grown in a growth chamber were measured. For quantification of pavement cell bulge/interdigitation, single layer scanned images were overlapped by LAS AF Lite software and then analyzed by Photoshop CS6 software. A stereomicroscope (Leica S8 APO and Leica DM6000B) was used to observe the sexual growth phenotype including flower, silique, stigma, and stamen. ImageJ was used to process the length measurement.

### Propidium Iodide (PI) Staining

For, PI staining, 10 μg/mL Propidium iodide (Sigma) was used to stain pavement cells for two days.

### Microscopy

For PI signal detection, a Leica TCS SP8 SMD microscope system was used. The argon laser was switched to 488 nm to detect GFP signal between 500 nm and 550 nm. The argon power was set at 20%. While 561 nm argon laser with power of 5% was used to detect PI signal between 560 nm and 580 nm.

## ACKNOWLEDGMENTS

No conflict of interest was declared. We would like to thank ABRC for providing seeds of SALK_016360 and SAIL_1242_H07, and to thank the NASC for providing seeds of SALK_129759 and GABI_191D02. This study was supported by National Natural Science Foundation of China (Grant 31470359) to J. H.

## ACCESSION NUMBER

Sequence data from this study can be found in the *Arabidopsis* Genome Initiative or GenBank/EMBL databases under the following accession numbers *TMK1* (AT1G66150), *TMK2* (AT1G24650), *TMK3* (AT2G01820), *TMK4* (AT3G23750).

## SUPPORTING INFORMATION

**Supplemental Table 1.**
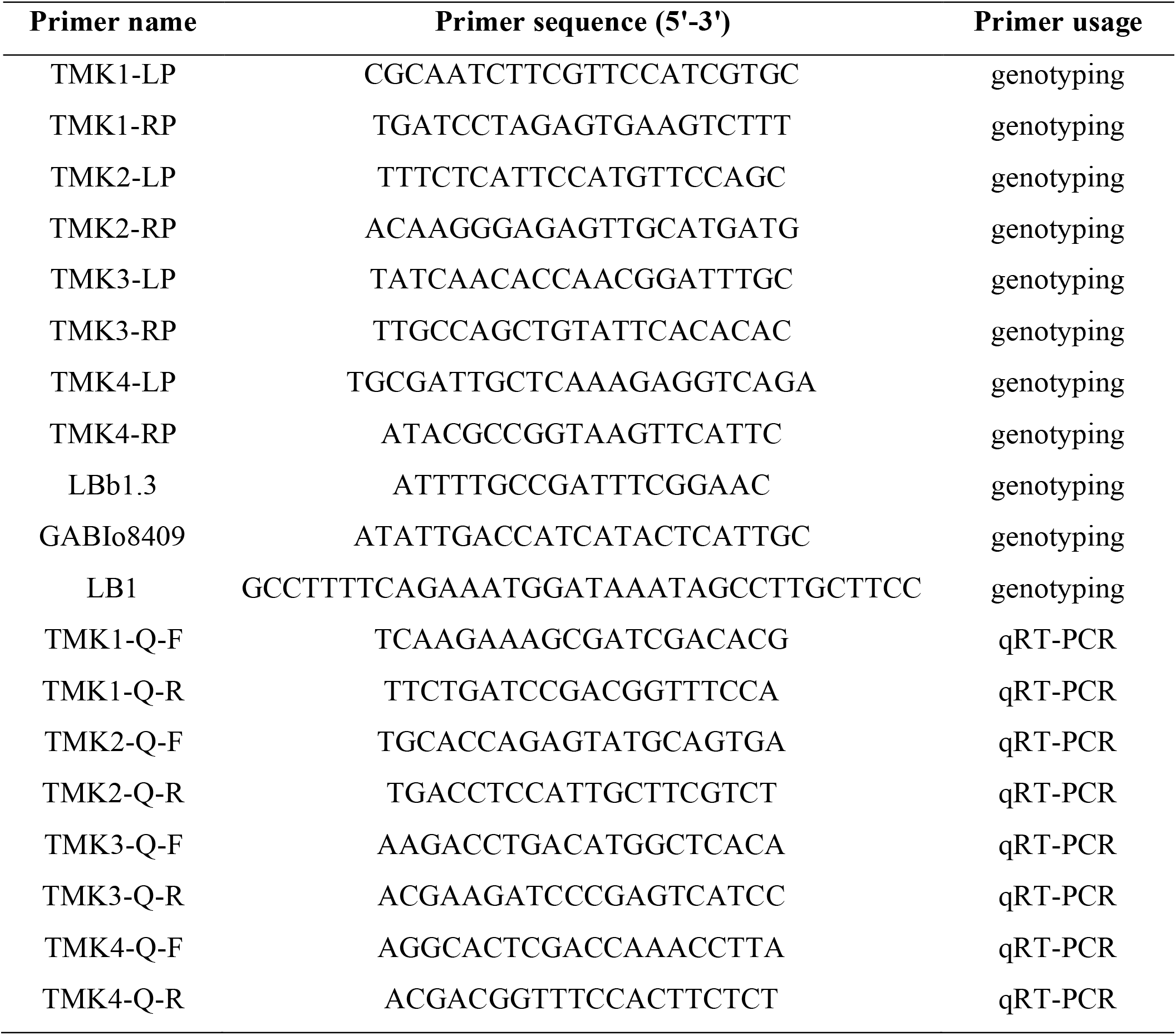
List of primers used in this study.

## Notes

Financial source: This work was supported by National Natural Science Foundation of China (Grant 31470359) to J. H.

### Competing Interest Statement

The authors have declared no competing interest.

